# GEM-based metabolic profiling for Human Bone Osteosarcoma under different glucose and glutamine availability

**DOI:** 10.1101/2020.09.08.287342

**Authors:** Ewelina Weglarz-Tomczak, Demi J. Rijlaarsdam, Jakub M. Tomczak, Stanley Brul

**Author notes:** shared first authorship (Ewelina Weglarz-Tomczak, Demi J. Rijlaarsdam).

## Abstract

Cancer cell metabolism is dependent on cell-intrinsic factors like genetics, and cell-extrinsic factors like nutrient availability. In this context, understanding how these two aspects interact and how diet influences cellular metabolism is important for developing personalized treatment. In order to achieve this goal, genome-scale metabolic models (GEMs) are used, however, genetics and nutrient availability are rarely considered together. Here, we propose an integrated metabolic profiling, a framework that allows to enrich GEMs with metabolic gene expression data and information about nutrients. First, the RNA-seq is converted into Reaction Activity Score (RAS) to further scale reaction bounds. Second, nutrient availability is converted to Maximal Uptake Rate (MUR) to modify exchange reactions in a GEM. We applied our framework to the human osteosarcoma cell line (U2OS). Osteosarcoma is a common and primary malignant form of bone cancer with poor prognosis, and, as indicated in our study, a glutamine-dependent type of cancer.

## Introduction

A widely recognized hallmark of cancer is reprogramming of cellular metabolism which aims at promoting the rapid cell proliferation and long-term maintenance, thus facilitating the uptake and conversion of nutrients into biomass^1–6^. Determining the metabolic requirement of proliferating cancer cells in order to modulate their metabolism might be a key factor to improve cancer treatment^7^. The genetic alterations in oncogenes and tumor suppressor genes promote cancer reprogramme cellular metabolism supporting tumorigenesis^1,2,8^. Therefore, the energy and nutrient requirements necessary for rapid proliferation can be realized. However, genetic alteration is not the only cell-intrinsic determinant of cancer cell metabolism; the second is the origin of the cell. All cancers begin when one or more genes in a cell mutate^9^. Therefore, there is not one cancer-type cell, and each cancer cell combines the metabolic features that come from the origin tissue/organ and from the mutation(s).
Understanding the regulation of the metabolic pathways by genetic factors provide new insights into the required metabolic aspect of tumorigenesis and can also potentially provide therapeutic metabolic targets.
Cancer cell metabolism similar to any other cell in living organisms is also determined by cell-extrinsic factors, including interactions with the environment^7,10^. There are numerous environmental factors that can affect cancer cell metabolism, where in particular diet, which affects nutrient availability, is one of the most important determinants^1,2,7,11,12^.
Many metabolic signatures are similar across different kinds of cancer cells like the Warburg Effect, where cells prefer aerobic glycolysis over oxidative phosphorylation followed by lactic acid fermentation, even in the presence of oxygen and functioning mitochondria^3,6^. However, it does not mean that all cancer cells are addicted to glucose and can not survive without it.

Here, we aim to contribute to the recently growing understanding of how certain cancer types respond to various nutrient availability and how the access to and utilization of nutrients by cancer cells affect the rate of proliferation. We provide a tool based on a genome-scale metabolic model (GEM)^13–17^ for approaching and testing hypotheses on how nutrient availability affects cancer cell metabolism and progression.

Furthermore, opposing the modulation of genetics, modulating the access to nutrients seems to be more feasible and safe. However, first the metabolic requirements of a particular cancer type should be defined.

In order to determine the nutrient requirement, we propose to utilize GEMs, transcriptomic and nutrient availability data. Moreover, we investigate nutrient requirements of cancer cells in different conditions with altered availability of nutrients, and examine its influence on transcription of metabolic genes.

In this paper, we focused on the human osteosarcoma cell line (U2OS). Osteosarcoma is a common and primary malignant form of bone cancer. It generally affects children and adolescents, where it represents the eighth-most common form of childhood cancer. With its poor prognosis, it is the second most important cause of deaths related to cancer in both children and adolescents^18^.

This paper has a multidisciplinary character, ranging from genomics and metabolism to *in silico* modeling. Therefore, we state the following research questions:

1. Do nutrient availability changes affect metabolism of the U2OS cells?
2. Do changes in glucose and/or glutamine availability impact the growth rate and metabolic requirement of U2OS cells?
3. Does simultaneous integration of the gene expression and nutrient availability data into a GEM provide accurate prediction of cell growth and metabolite consumption/production?

To achieve our goals we performed integrated metabolic profiling of the U2OS cell line that includes experiments and genome scale modeling. By gathering transcriptomic data from these cells in varying microenvironments, the effect of nutrient availability on cancer cell metabolism was studied. We focused on the questions whether this cell line is glucose-dependent, and whether U2OS cells require glutamine in order to grow and survive. Finally, we addressed the question whether diet (using altered nutrient availability) affects the transcription level of the cells and can alter the expression of the metabolic profile. As a modeling platform we used the genome-scale metabolic models^13–17^ that can be used to predict metabolic flux values for metabolic reactions using optimization techniques, such as Flux Balance Analysis (FBA)^16,17^ and Flux Variability analysis (FVA)^19^, which uses linear programming. GEMs allow to describe an entire set of stoichiometry-based, mass-balanced metabolic reactions with gene-protein-reaction (GPR) associations.

In our study, we utilized the most comprehensive human metabolic network model to date, Recon3D^20^, and a novel method, named *Gene Expression and Nutrients Simultaneous Integration* (GENSI)^21^ to create cell specific models. GENSI translates the relative importance of gene expression and nutrient availability into the fluxes. Further, we used our models to predict the influence of glucose and glutamine availability on the biomass synthesis flux and cancer related metabolic behaviors like the consumption of glucose and glutamine and the production of lactate. Our integrated study that includes experiments on the cell line and the computational simulation on RECON3D-based metabolic reconstruction shows that the U2OS is independent of the glucose level and the presence of glutamine is essential for cell viability.

The contribution of this paper is sixfold:

1. We provide an integrated GEM-based metabolic profiling framework for identification of the nutrient requirements.
2. We prove that genome-scale models like RECON3D provide a great platform for integration of transcript and nutrient availability data in order to create specific models for studying cell metabolism.
3. We prove that translation of the relative importance of gene expression and nutrient availability data into the fluxes based on observed experimental feature(s) is a reliable method to build specific models.
4. We identified the impact of glucose and glutamine availability on the metabolism and proliferation rate of the U2OS cells.
5. We identified that the U2OS cell line is a glutamine-dependent cancer type.

## Results

### GEM-based metabolism profiling methodology

To study the influence of nutrient availability on the growth rate and the metabolic behaviour we used the U2OS cell line and seven DMEM derived media containing different concentrations of glucose and glutamine (or *L*-alanine-glutamine), see Table 1 in the Materials & Methods section. The U2OS cell line consists of epithelial adherent cells and exhibits a fast growth rate.

**Table 1.** The seven different medium compositions used to culture U2OS cells.

| Condition name | Glucose (mM) | Glutamine (mM) |
| --- | --- | --- |
| Glc10Gln4 | 10 | 4 |
| Glc2Gln4 | 2 | 4 |
| Glc10Gln0 | 10 | 0 |
| Glc2Gln0 | 2 | 0 |
| Glc25Gln4 | 25 | 4* |
| Glc2Gln0.5 | 2 | 0.5 |
| Glc2Gln2 | 2 | 2 |

\*L-Alanyl-Glutamine

We cultured them altering the metabolic microenvironment until the growth rate achieved a steady state. Further, we performed the proliferation assays and isolated mRNA from the cells and metabolites from the medium. We performed RNA-seq analysis using the NextSeq 550 Sequencing System (Illumina) and generated transcriptomic data after proper quality control and filtration. The RNA sequencing data were then processed (see Materials & Methods) and used for creation of the specific GEM models. Since GEMs include known functions of protein-encoding genes, they can be used as platforms for analyzing mRNA expression data to elucidate how changes in gene expression impacts cell metabolism and phenotypes^22–28^. Here, we additionally integrated the nutrient availability data by constraining the bounds of the exchange reactions. Our supervised approach that integrates both RNA-seq and nutrient availability data is based on the recently published GENSI^20^ method that utilizes the *Reaction Activity Score* (RAS) approach for mapping the transcriptome into a metabolic network^23^. GENSI creates GEM-based specific models by integrating both data into the fluxes based on observed feature(s) that distinguish conditions. The proposed integrated metabolic profiling framework is schematically depicted in Fig. 1.

**Fig. 1.**
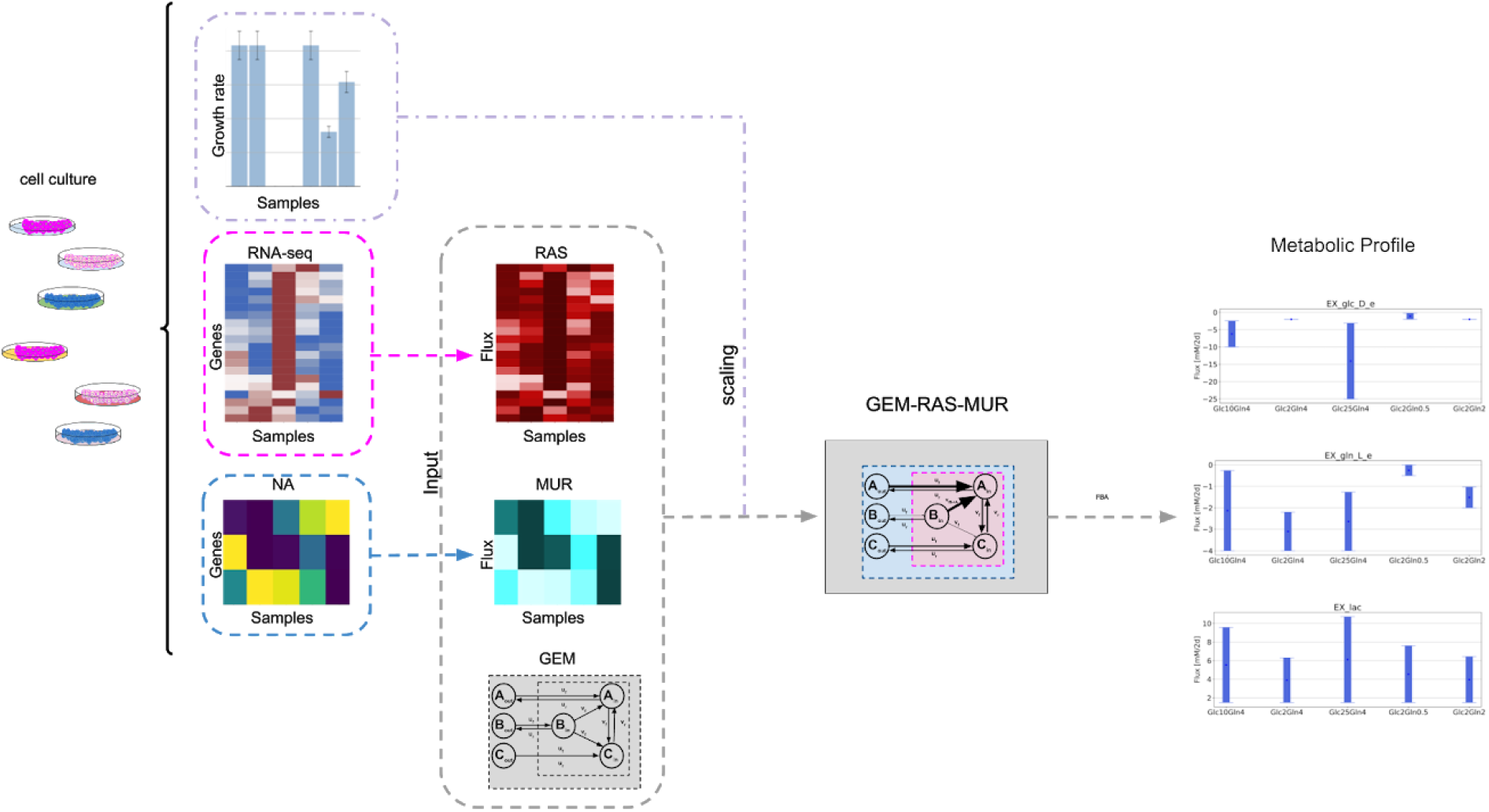
A visualization of the steps in GEM-based metabolic profiling. GEM-based integrated metabolic profiling requires three inputs: a GEM model, gene expression and nutrient availability data and one feature of the cells that distinguish cells cultured in different conditions. In the first step of GENSI^21^, the RNA-seq is converted into RAS score^24^ that is defined for any reaction *i* associated with a gene in the GEM as a function of the expression of the genes encoding for the *subunits* and/or the *isoforms* of the associated enzyme(s). Nutrient Availability (NA) is converted to Maximal Uptake Rate (MUR) that is defined for an exchange reaction *j* in GEM and describes the rate of the maximum possible uptake over the time for substances available for cells. NA is limited by a composition of media used in our experiments. Obtained GEM-RAS-MUR models are then used for metabolic profiling using optimisation methods such as FBA^16,17^ and FVA^19^.

### Formulation of the specific GEM model

A central challenge in understanding and treating cancer comes from its multi-scale nature. To gain an insight into the impact of the access to nutrients on metabolism and the rate of biomass synthesis, we integrated both; transcriptomic and nutrient availability data into the most extensive genome-scale model of human metabolism RECON3D^21^ and studied their meaning using the FBA^16,17,19^ approach.

In the GENSI framework the flux bounds of the reactions that are associated with genes are set proportionally to *RAS* scores that are computed based on RNA-seq data. In our work, the flux bound *v_i_* of the reaction *i* is equal to RAS*_i_* multiplied by a factor α (*v_i_* = α × *RAS_i_*), where α is the factor independent of the reaction identity. When reaction *i* is reversible, its forward flux is bounded by *v_i_* and its backward flux is bounded by −*v_i_*. When the reaction *i* is irreversible, the flux bound in the impossible direction remains zero and the flux bound in the possible direction is set to *v_i_*. While exchange fluxes are correlated with relative estimates of consumption and/or secretion rates. Nutrient availability data was converted into a maximum uptake rate that describes the rate of the maximum possible uptake over the time for substances available for the model. Here, we defined *MUR_j_* for the exchange reaction *j* in the GEM as the absolute value of a difference in concentration of the substance *s_j_* in an extracellular environment over 48h, for more details see Materials & Methods.

Following the GENSI framework, we found the value of the factor α by computing the steady state flux pattern for the maximal biomass synthesis flux for various combinations of RAS and MUR within GEMs. Possible values of α were between 0.000001 and 1, see Fig. 2. For the factor value equal to 0.00042, we observed that predictions for limitations imposed by nutrient availability and gene expression, i.e., GEM-RAS-MUR, matched almost ideally maximal biomass synthesis fluxes corresponding to the proliferation rate observed in experiments (Fig. 2c and Fig. 3c). Interestingly, the growth rate does not depend on the glucose concentration in the presence of 4 mM of the glutamine (or *L*-alanine-glutamine).

**Fig. 2.**
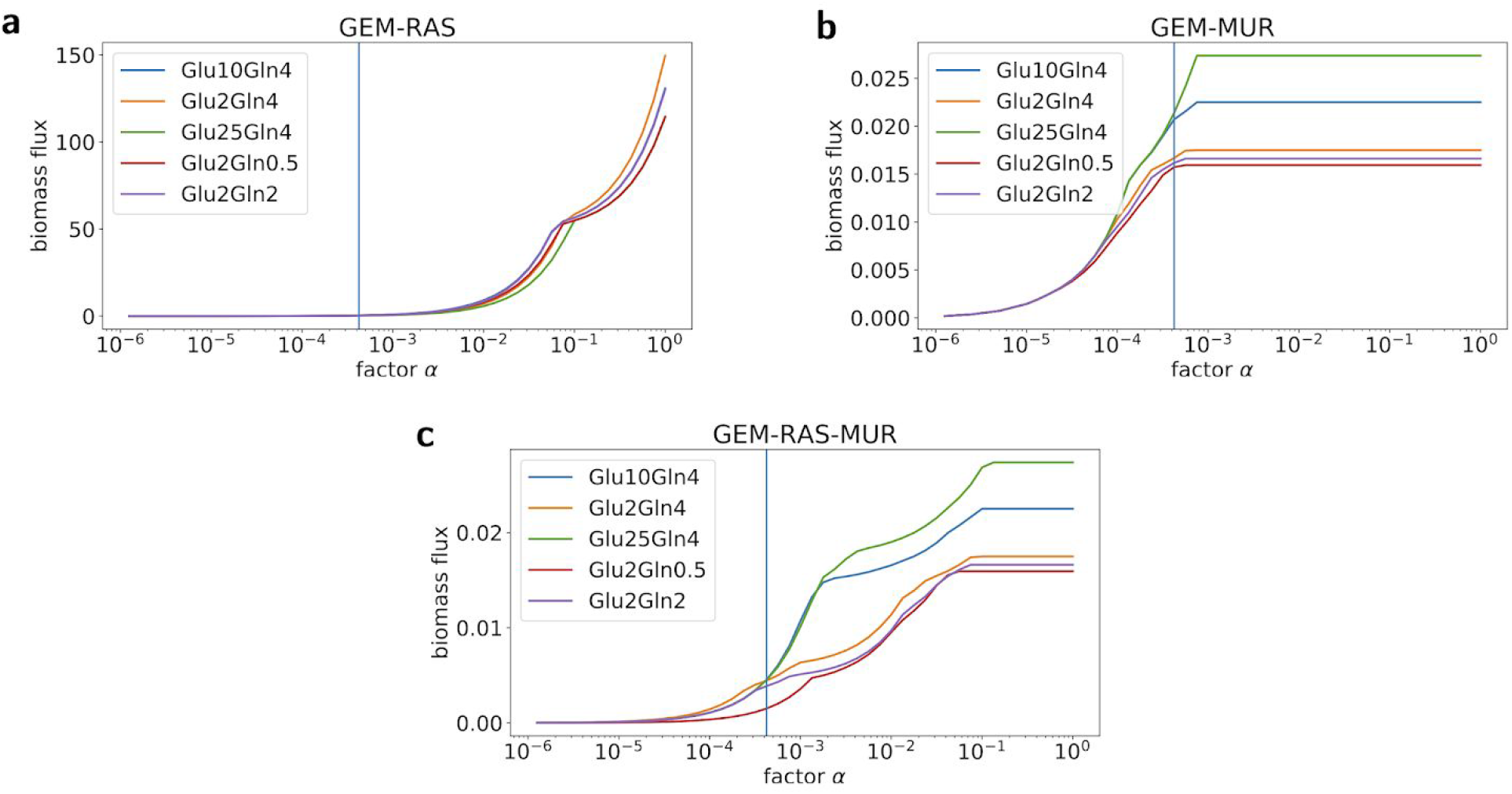
Dependencies between values of in-silico biomass flux (y-axis) and values of factor α (x-axis) for different combinations of GEMs with RAS and MUR. **a** The dependency for GEM with RAS and without MUR. **b** The dependency for GEM with MUR and without RAS. **c** The dependency for GEM with RAS and MUR.

**Fig. 3.**
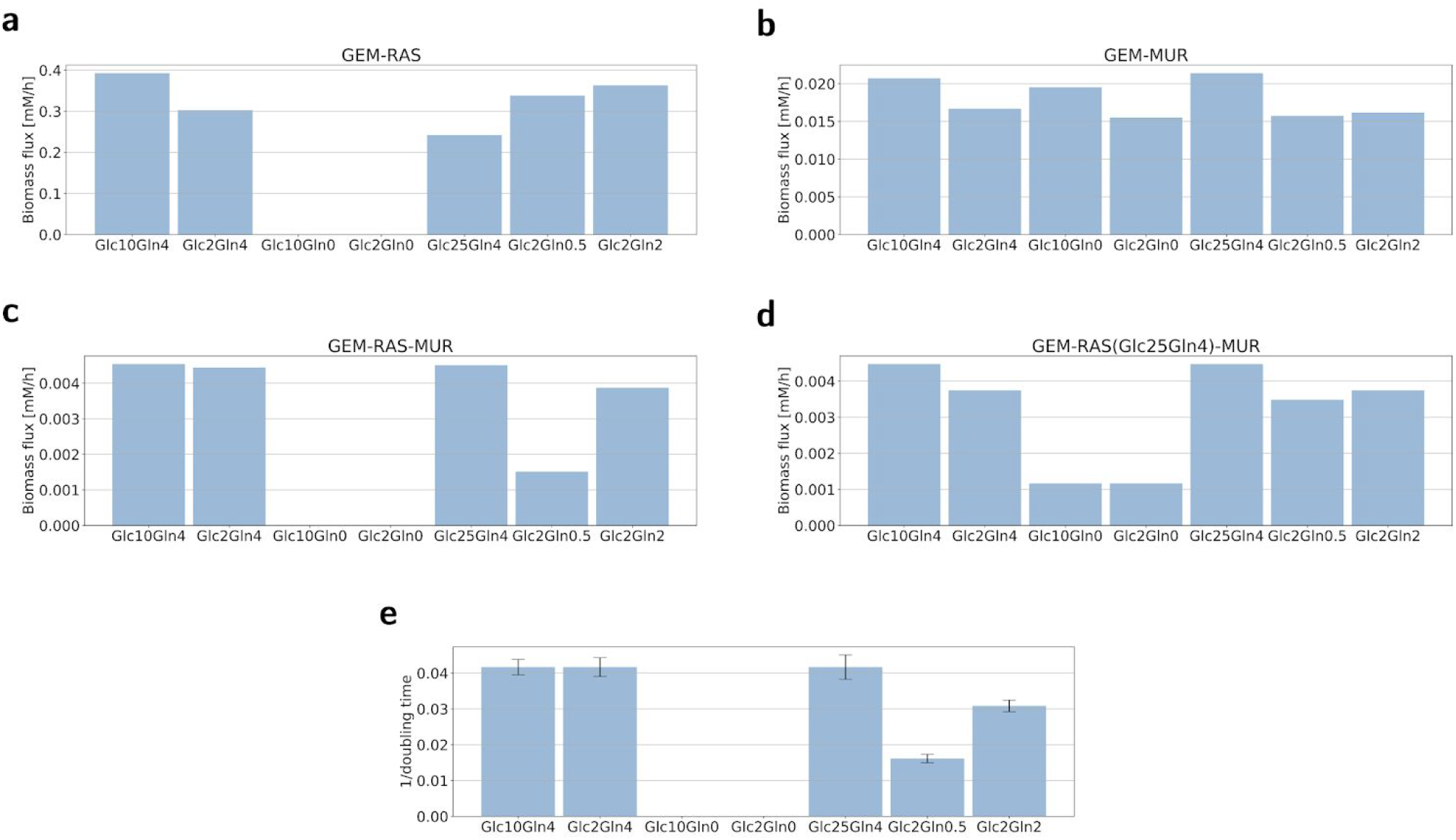
*In-silico* biomass flux predictions and observed proliferation growth rates. **a,b,c,d** Maximal *in-silico* biomass flux predictions for U2OS across the specific models created for each condition based on: transcriptomic data (GEM-RAS)* (**a**), nutrients availability data (GEM-MUR)* (**b**), transcriptomic data and nutrient availability data (GEM-RAS-MUR)* (**c**), transcriptomic data from the condition Glc25Gln4 and nutrient availability data (GEM-RAS(Glc25Gln4)-MUR) (**d**). **e** Reciprocal of the cells doubling time calculated from a linear equation describing the change in cell number over the time. ** * No data for conditions Glc10Gln0 and Glc2Gln0 due to the fact that cells were not able to survive and the RNA could not be isolated. ** For Glc10Gln0 and Glc2Gln0 the doubling time was not available due to the fact that the growth rate was less than or equal to zero.

In the absence of glutamine, the cells were unable to survive and started dying after hours of culturing (Fig. 3e). Since cells could not proliferate without glutamine we were not able to create GEMs for Glc10Gln0 and Glc2Gln0 due to lack of the RNA-seq data. We then built GEM-RAS(*Glc25Gln4*)-MUR models using RNA-seq data from cells cultured in Glc25Gln4 medium, which was our starting condition.Computed biomass fluxes across seven different nutrient availability data (Fig. 3d) clearly show that without glutamine the U2OS cells have the least chance of survival. The biomass flux for the cases without glutamine was much smaller (Fig. 3d). The results of this prediction are consistent with the viability of cells and show that the presented framework is a good approach to predict survival of cells in new conditions.

To examine whether only gene expression level has impact on the biomass synthesis, we formulated five RECON3D-based models via constraining reactions associated with genes as a function of RAS without changing exchange reactions (GEM-RAS). We did observe the impact of mapping the gene expression, however, there was no value of the factor α for which predicted biomass fluxes and growth rates are correlated (Fig. 3a). This result seems to confirm that genetics is not the only determinant of the metabolic phenotype.

Next, we decided to test whether integration of only nutrient availability changes the prediction of the fluxes (GEM-MUR). We built seven GEM-MUR models via constraining exchange reactions by MUR calculated for each condition. We did not change the bounds of reactions associated with genes. As we expected, the results again did not meet the experimental observations, leading even to an experimentally unobserved situation of a high growth rate for Glc10Gln0 and Glc2Gln0 (Fig. 3b).

The predictions of biomass flux for GEM-RAS-MUR models were aligned with our proliferation assay (compare Fig. 3c and Fig. 3e); the biomass flux proportional to the reciprocal doubling time is glucose-independent and decreases as the amount of glutamine changes.

### Flux variability analysis to investigate the nutrient requirement

To test the prediction of the GEM-RAS-MUR models we performed FVA analysis for allowed exchange reactions while supporting biomass production rate. We further explored the range of uptake (if negative) or secretion (if positive) fluxes of glucose, glutamine and lactate by plotting the range of fluxes for corresponding exchange reactions (Fig.4).

Our main observation was that glutamine uptake is essential for all conditions. The minimum of the flux of the glutamine exchange reaction was alway less than zero (Fig. 4c) (however, for the condition Glc2Gln0.5 the value was relatively small). In medium Glc2Gln0.5 it is then required at a very low amount of glutamine to achieve optimal growth. Whereas the minimum uptake of glucose is much less than its availability and equal to 2mM/48h for most conditions (Fig. 4a). Only for the condition with Glc2Gln0.5 the minimum flux was equal to 0.24 mM/48h. The overall prediction of the maximum flux of lactate seems to be in accordance with the glucose consumption: around 10mM/48h for conditions Glc25Gln4 and Glc10Gln4 and 7mM/48 for conditions containing 2mM of glucose (Fig. 4e).

**Fig. 4.**
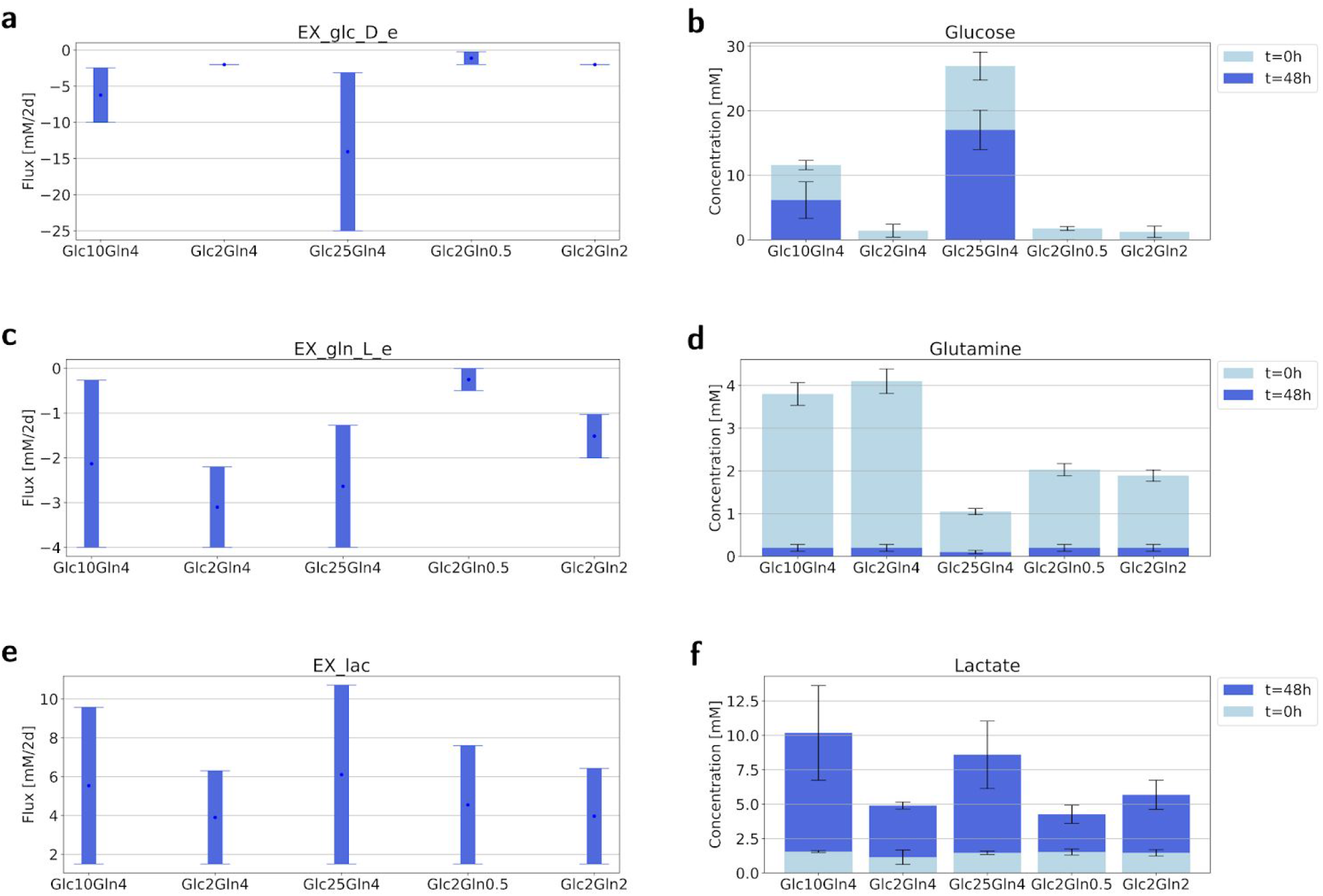
Predicted and observed ranges of the uptake/secretion of glucose, glutamine and lactate. **a**,**c**,**e** The range of fluxes of glucose (**a**), glutamine (**c**) and lactate exchange reaction (**e**) consistent with the maximal biomass flux and steady state across five different media. **b**,**d**,**f** The concentration of glucose (**b**), glutamine (**d**) and lactate (**f**) in five different media before and after carrying out experiments.* * The concentration of the cells at t=0 was 200 000 cells/ml.

*In-silico* predictions were then verified experimentally by measurement of the glucose and glutamine consumption, and lactate consumption. It turned out that all of the available glutamine were consumed by cells in all cases. While the excess of glucose in the conditions Glc10Gln4 and Glc25Gln4 were not utilized by cells. Interestingly, the production of lactate was similar for Glc10Gln4 and Glc25Gln4, while cells cultured in Glc25Gln4 consumed two times more glucose than cells growing in Glc10Gln4.

## Discussion

Cancer is one of the world’s most serious health problems. Recently, cancer metabolism and nutrition have gradually become important topics in theoretical and clinical research. In contrast to traditional chemotherapy, where the goal is to kill cancer cells mainly via blocking the transcription, the nutrition-based approach assumes that cancer cell metabolism is influenced by metabolic constraints imposed by a diet. Perturbations to dietary compositions contribute to changes in plasma metabolite levels that in turn influence metabolite levels in the tumor microenvironment and thereby alter cancer cell metabolism and therapeutic responses. Nutrient levels in the tumor/cancer cell microenvironment can have profound impacts on the cell metabolism, growth and drug sensitivity^7,10–12^.

Here, we follow this line of research and propose a framework for exploring the impact of the diet on cancer cell metabolism and progression using the cancer cell line and medium compositions as an experimental setup. We performed the integrated metabolic profiling through enriching genome-scale modeling with metabolic gene expression and nutrient availability data. In our study, we focused on the metabolism of bone osteosarcoma that is one of the most common primary malignant bone tumors^18^.

As a GEM model we used Recon3D^20^ that is a computational resource and functions as the most extensive, updated and expanded human metabolic network model. With its efficiency to connect genes to biochemical pathways, it can emphasize the promising prospects of structural analysis in genome-scale models for identifying genes related to disease and the potential for developing new treatment, biomarkers or drug repurposing. Moreover, as we recently showed, it could also be utilized as a platform for integration of the cell-intrinsic factor (gene expression level) and cell-extrinsic factor (nutrient availability) and contribute to the understanding of the impact of the nutrients availability on cancer metabolism^20^.

Here, we proposed a general framework of the integrated metabolic profiling that incorporates information about gene expression and nutrient availability to a GEM that could be further used to accurately predict cell growth rates and nutrients/metabolites consumption/production. For this purpose, we utilized the GENSI framework to scale reaction bounds using RAS, and then MUR to introduce information about nutrient availability to external reactions.

In our experiments, we considered seven different conditions in which various amounts of glucose and glutamine (mM) were available, see Table 1 in Materials & Methods. First, we noticed that in the case of no glutamine (Glc10Gln0 & Glc2Gln0), all cells died within two weeks; that was a clear indication of the importance of the glutamine availability for the cell growth. Furthermore, the conditions Glc10Gln4 and Glc2Gln4, as well as the condition Glc25Gln4, displayed roughly similar and high growth rates (Fig. 3e), indicating a more important role of glutamine compared to glucose in the growth of U2OS cells. This fact is further supported by the growth rates of the conditions Glc2Gln0.5 and Glc2Gln2, where Glc2Gln2 clearly presented a higher growth rate (almost as high as conditions with Gln4) in comparison to Glc2Gln0.5. We were able to answer our research questions (1 and 2) that the U2OS cell line is a glutamine-dependent cancer type and glucose has very limited influence on the cancer cell growth.

Next, we showed that our integrated metabolic profiling allowed us to obtain a GEM that provides highly accurate predictions. For this purpose, we first verified whether a combination of RAS and MUR (GEM-RAS-MUR) is crucial to match model predictions with observed quantities and compared them against a GEM with information about RAS (GEM-RAS) and a GEM with MUR (GEM-MUR). The results in Fig. 3 seem to confirm that genetics or nutrient availability used separately are insufficient determinants of the metabolic phenotype. Only their combination (GEM-RAS-MUR) allowed us to obtain accurate predictions of the cell growth.

Lastly, we applied flux variability analysis to the GEM obtained within the proposed framework (GEM-RAS-MUR) to compare uptake/secretion fluxes of selected metabolites with measured quantities. In Fig. 4 we confronted predicted fluxes by the FVA (Fig. 4a, 4c & 4e) against the measured quantities (Fig. 4b, 4d & 4f) of glucose, glutamine and lactate. Interestingly, the predictions given by the FVA closely reflect real consumption/production. This outcome provided further evidence that the proposed integrated metabolic profiling framework can be used to develop models capable of accurately imitating highly complex processes like cancer metabolism.

In conclusion, the results obtained in our experiments indicate that the GEM-RAS-MUR model for the U2OS cell line obtained within our framework could be used to accurately predict cell growth and metabolite consumption/production. This answers our last research question positively and opens new research perspectives for applying our framework to different types of cancer, but also other illnesses. The combination of transcriptomic data together with information about nutrients seems to be essential for developing accurate *in-silico* models like GEMs. Lastly, our study also highlights the huge potential of GEMs as a platform to incorporate big data like RNA-seq.

## Materials & Methods

### Culturing U2OS Cancer Cells

U2OS (Human Osteosarcoma) cells were cultured in Dulbecco’s Modified Eagle Medium (DMEM; GibcoBRL, Grand Island, NY, USA), supplemented 10% Fetal Bovine Serum (FBS) and 1% antibiotics (penicillin and streptomycin) at various initial concentrations of glucose and glutamine/L-alanine-glutamine (Table 1). Six of the compositions were made using Advanced DMEM that does not contain glucose, glutamine, phenol red and the sodium pyruvate. Glc25Gln4 is commercially available (DMEM glutamax™). The cells were first cultured in DMEM glutamax™ for two weeks until a sufficient amount of cells was acquired. Then cells were split over seven experiments and cultured for three weeks to adopt. Cells cultured in condition 3 and 4 started dying immediately.

### Growth rate/Cell proliferation assay

The total number of cells in the consequent supernatant was determined by hemocytometer counting. Mean growth rate was determined by counting cells in four non-overlapping sets of sixteen corner squares selected at random.

### Isolation and measurement of the metabolites from medium

Samples from each medium were collected in time 0 and 48h, repeated three times for a total of three independent experiments for every medium composition. Metabolites were isolated using the protocol described in Sapcariu *et al*^29^ with minor modification. The samples were filtered using 0.22μm syringe filters (BGB Analytik Vertrieb GmbH, Rheinfelden, Germany), freeze dried and stored until measurement in −80°C. Standard solutions for glutamine, glucose and lactate were prepared in the same way as well as control samples. *High-performance liquid chromatography* (HPLC) was performed using the HPLC-DAD_RID LC-20AT Prominence (Shimadzu, Columbia, USA) machine with a UV Diode Array Detector SPD-M30A NexeraX2 or/and a Refractive Index Detector RID 20A and an analytical ion-exclusion Rezex ROA-Organic Acid H+(8%) column (250×4.6 mm) with guard column (Phenomenex, Torrance, USA) (5 mM H2SO4 in MilliQ water (18.2 MΩ),isocratic, 0.15 ml/min. flow rate). Injection volume was 15 μl (Autosampler: SIL-20AC, Prominence, Shimadzu), column oven temperature was 55 °C (Column oven: CTO-20A, Prominence, Shimadzu) and the pressure was 29 bar.

### Isolation and analysis of RNA

5.000.000 cells from each medium condition were taken for RNA analysis, also repeated three times for a total of three independent experiments for every cell condition. The RNA extraction was done using the TRI Reagent Protocol from Sigma Aldrich (https://www.sigmaaldrich.com/technical-documents/protocols/biology/tri-reagent.html), after which all samples were treated with DNase using the TURBO DNA-free Kit from Invitrogen. Nanodrop was used for determining RNA concentrations when needed. With the Agilent 2100 Bioanalyzer System the quality of the RNA was checked, as well as the RNA Integrity Number (RIN), using RNA Pico Chips. rRNA depletion and library preparation for sequencing was completed with the NEBNext protocol (‘Protocol for use with NEBNext rRNA Depletion Kit (Human/Mouse/Rat) (NEB #E6310) and NEBNext Ultra II Directional RNA Library Prep Kit for Illumina (NEB #E7760, #E7765)’) from New England BioLabs. Using a 2200 TapeStation System with Agilent D1000 ScreenTapes (Agilent Technologies), the size distribution of the libraries with indexed adapters was assessed. Quantification of the libraries was performed on a QuantStudio 3 Real-Time PCR System (Thermo Fisher Scientific), which was done using the NEBNext Library Quant Kit for Illumina (New England BioLabs) according to the manufacturer’s directions. On a NextSeq 550 Sequencing System (Illumina), the libraries were clustered and sequenced (75 bp) utilizing a NextSeq 500/550 High Output Kit v2.5 (75 Cycles) (Illumina).

### RNA-seq data processing

RNA-seq data was available in .fastq file format. There were three measurements per condition. The RNA-seq data was processed according to the following pipeline:

1. Sequences were mapped against the human reference genome using Burrows-Wheeler Aligner (BWA; version 0.7.17, https://sourceforge.net/projects/bio-bwa/). The output of this step were files in .sam format.
2. The .sam files were transformed to sorted bam files using Samtools (https://anaconda.org/bioconda/samtools).
3. Gene occurrences were calculated using featuresCounts from the Subread package (https://anaconda.org/bioconda/subread).

In all cases, the *Homo Sapiens* (human) genome assembly GRCh38 (hg38) from Genome Reference Consortium Build 38 (https://www.ncbi.nlm.nih.gov/assembly/GCF_000001405.26/) was used as the reference genome.

### Mapping transcriptomic data to the fluxes with Recon3D

Mapping transcriptomics data to the fluxes with Recon3D includes two steps: 1) Conversion of RNA-seq into RAS, 2) RAS to fluxes of reactions with Gene-Protein-Reaction association rules.

RNA-seq data was converted to RAS, for each condition and each reaction, based on protocol declined by Graudenzi, A. et al.^24^ according to Gene-Protein-Reaction association rules (GPRs). The formula includes *AND* and *OR* logical operators. AND rules are employed when distinct genes encode different subunits of the same enzyme, i.e., all the subunits are necessary for the reaction to occur. OR rules describe the scenario in which distinct genes encode isoforms of the same enzyme, i.e., either isoform is sufficient to catalyze the reaction. In the next step *RAS*_i_ is converted to a lower bound *v_i_* of the reaction *i* having GPR association rules as follows:

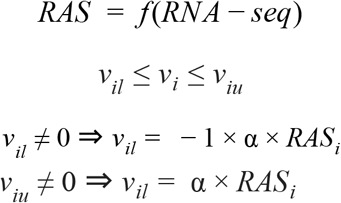

where *v* is the vector of fluxes through all reactions having a gene rule, *v_il_* and *v_iu_* are the vectors of lower and upper bounds on the reaction *i*.

### Mapping Nutrient Availability (NA) to fluxes with Recon3D

Mapping NA to fluxes with Recon3D includes two steps: 1) Conversion of NA to Maximum Uptake Rate (MUR), 2) Conversion of MUR to fluxes of exchange reactions. We define a *MUR_j_* for each exchange reaction *j* based on NA data for each condition. In the next step *MUR_j_* is converted to a lower bound *v_jl_* of exchange reaction *i* as follows:

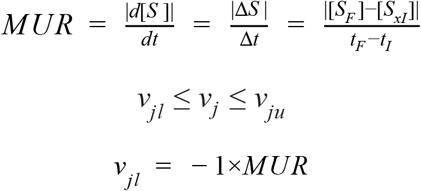

where *v_i_* is the vector of fluxes through exchange reactions with the environment of the system, *v_jl_* and *v_ju_* are the vectors of lower and upper bounds on these fluxes, *MUR* is a Maximum Uptake Rate that is defined as the rate of the maximum possible uptake over the time for each substance available to the model.

In our case, MUR is defined for the exchange reaction *j* in a GEM as the absolute value of the difference in the concentration of the substrate *S_j_* in extracellular environment (|[*S_jF_*] − [*S_jI_*] |) over the 48h.

### Simulations

In the simulations of maximal biomass flux we applied the computational technique of Flux Balance Analysis (FBA)^16^ to the human genome-wide metabolic map Recon3D^20^ using the COBRA toolbox in MATLAB and Python^30,31^. FBA solves the following linear programming problem:

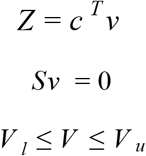

where *S* is the stoichiometry matrix indicating how many molecules of each metabolite are produced or consumed in each reaction, *V* is the vector of fluxes through all reactions including exchange reactions with the environment of the system, *V_l_ a* and *V_u_* are the vectors of lower and upper bounds on these fluxes, ***c*** is a vector of weights generating the linear combination of fluxes that constitutes the objective function ***Z***.

In the simulations of maximal and minimal fluxes for defined exchange reactions we applied the computational technique of Flux Variability Analysis (FVA)^19^. FVA is used to find the minimum and maximum flux for reactions in the network while maintaining some state of the network, e.g., supporting maximal possible biomass production rate. FVA entails the following linear programming problem:

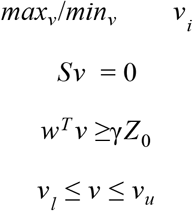

where *Z*_0_ = *w*^*T*^*v*_0_ is the optimal solution to the FBA problem with biomass reaction as the objective function, *w^T^* represents the biomass objective (vector of weights generating the linear combination of fluxes that constitutes the objective function *Z*), *v* is the vector of fluxes through all reactions, γ is a parameter that controls whether the analysis is done w.r.t. suboptimal network states (0≤γ <1) or to the optimal state (γ = 1).

## Data and source code availability

All related data sources such as gene featureCounts, RAS scores, and GEM models can be found at https://github.com/e-weglarz-tomczak/GEM-RAS-MUR

## Acknowledgments

We gratefully acknowledge Will Beckam for his kind help with RNA isolation and Eugenie Troia for her help with HPLC analysis.

EWT is co-financed by a grant Mobilność Plus V from the Polish Ministry of Science and Higher Education (Grant No. 1639/MOB/V/2017/0).

## Author contributions

EWT conceived the project with advice from JT and SB. DR performed all *in vitro* experimental work with EWT supervision. JT processed gene expression data. EWT prepared GEMs and performed flux-based analyses. DR, EWT, JT and SB drafted and revised the manuscript.

## Materials and correspondence

Correspondence and requests for materials should be addressed to EWT.

